# Both clinical and environmental *Caulobacter* species act as opportunistic pathogens

**DOI:** 10.1101/645515

**Authors:** Gabriel M. Moore, Zemer Gitai

## Abstract

The *Caulobacter* genus, including the widely-studied model organism *Caulobacter crescentus*, has been thought to be non-pathogenic and thus proposed as a bioengineering vector for various environmental remediation and medical purposes. However, *Caulobacter* species have been implicated as the causative agents of several hospital-acquired infections, raising the question of whether these clinical isolates represent an emerging pathogenic species or whether *Caulobacters* on whole possess previously-unappreciated virulence capability. Given the proposed environmental and medical applications for *C. crescentus*, understanding the potential pathogenicity and human health implications of this bacterium is crucial. Consequently, we sequenced a clinical *Caulobacter* isolate to determine if it has acquired novel virulence determinants. We found that the clinical isolate represents a new species, *Caulobacter mirare. C. mirare* phylogenetically resembles both *C. crescentus* and the related *C. segnis*, which was also thought to be non-pathogenic. The similarity to other *Caulobacters* and lack of obvious pathogenesis markers suggested that *C. mirare* is not unique amongst *Caulobacters* and that consequently other *Caulobacters* may also have the potential to be virulent. We tested this hypothesis by characterizing the ability of *Caulobacters* to infect the model animal host *Galleria mellonella*. In this context, two different lab strains of *C. crescentus* proved to be as pathogenic as *C. mirare*, while lab strains of *E. coli* were non-pathogenic. Further characterization showed that *Caulobacter* pathogenesis is mediated by a dose-dependent, cell-associated toxic factor that does not require active bacterial cells or host cellular innate immunity to elicit its toxic effects. Finally, we show that *C. crescentus* does not grow well in standard clinical culture conditions, suggesting that *Caulobacter* infections may be more common than generally appreciated but rarely cultured. Taken together, our findings redefine *Caulobacters* as opportunistic pathogens and highlight the importance of broadening our methods for identifying and characterizing pathogens.

**AUTHOR SUMMARY:** Bacterial species have historically been classified as either capable of causing disease in an animal (pathogenic) or not. *Caulobacter* species represent a class of bacteria that were thought to be non-pathogenic. *Caulobacters* have been widely studied and proposed to be used for various industrial and medical applications due to their presumed safety. However, recent reports of human *Caulobacter* infections raised the question of whether disease-causing *Caulobacters* have acquired special factors that help them cause disease or whether the ability to infect is a more general feature of most *Caulobacters*. By combining genomic sequencing and animal infection studies we show that a clinical *Caulobacter* strain is similar to lab *Caulobacters* and that all *Caulobacters* studied can infect a model host. We explore the mechanism of this infectivity and show that it is due to a toxic factor that associates with *Caulobacter* cells. We also provide a possible explanation for why *Caulobacters* have not traditionally been isolated from human patients, owing to their inability to tolerate the salt levels used in most medical culturing systems.

## INTRODUCTION

The free-living, gram-negative genus *Caulobacter* was first described and classified as a group of rod-shaped, stalk possessing bacteria in 1935 [1, 2]. Since their identification, *Caulobacter* have been observed in rhizosphere, soil, and aqueous environments, including drinking water reservoirs [3, 4]. Historically, this genus has been considered non-pathogenic due to lack of presence in infection cases, no obvious pathogenicity islands, and increased bacterial mortality at human body temperatures [5]. However, the last two decades has seen several reports of symptomatic infections associated with *Caulobacter* species [6–10]. All reported cases of *Caulobacter* infections appear to be hospital-acquired by immunocompromised patients, suggesting that these infections are opportunistic. None of the *Caulobacter* isolates associated with human infection have been previously sequenced. Consequently, it remains unclear whether clinical isolates have acquired virulence mechanisms absent from other *Caulobacters*, or if *Caulobacter* species generally have the capacity for human disease in the right context.

Among *Caulobacter* species, *Caulobacter crescentus* is the best characterized and most widely studied in laboratory settings [11]. *C. crescentus* has been primarily used as a model organism for understanding bacterial cell-cycle progression due to its highly regulated asymmetrical division and dimorphic lifestyle [12, 13]. Because of its available molecular tools, ability to display proteins in its surface layer (S-layer), and assumed non-toxicity to humans, *C. crescentus* has been proposed to be a powerful vector for a wide range of bioengineering applications [14, 15]. For example, *C. crescentus* has been engineered as a biosensor for uranium [16], a bioremediation tool for heavy metals [17], an anti-tumor immunization technique [18], and an anti-viral microbicide in humans [19]. Thus, understanding the potential pathogenicity and human health implications of this bacterium is crucial before its industrial use.

Here we obtained and sequenced a *Caulobacter* isolate from a reported human infection (7) to determine if it contains conspicuous virulence determinants or is similar to previously-characterized *Caulobacters*. We found that the clinical isolate represents a new species with similarities to both *C. crescentus* and *C. segnis*. The lack of pathogenicity islands and similarity to lab strains of *C. crescentus* suggested that the potential of this clinical isolate to be an opportunistic pathogen may be a general feature of *Caulobacters*. We confirmed this hypothesis by turning to the *Galleria mellonella* model animal host. The clinical *Caulobacter* isolate and lab strains of *C. crescentus* exhibited similar virulence, which were both significantly higher than non-pathogenic lab strains of *E. coli*. Further characterization revealed that *Caulobacter* pathogenicity is mediated by a toxic cell-associated factor. Our results thus redefine *Caulobacter* as a potential opportunistic pathogen and establish the interaction between *Caulobacter* and *G. mellonella* as a model for host-pathogen interactions.

## RESULTS

### Clinical *Caulobacter sp. SSI4214* shares homology with soil- and freshwater-associated species of *Caulobacter*

To genomically characterize a clinical *Cauloacter* isolate, we obtained a clinical strain of *Caulobacter* species isolated from the dialysis fluid of a 64-year-old man in Denmark with peritonitis [7]. There was only one bacterial species that could be cultured from the peritoneal fluid using Danish blood agar medium, and the infection responded to gentamycin treatment suggesting that this species was the likely cause of the infection [7]. Imaging of the cultured bacteria revealed a crescent-shaped morphology similar to that of *Caulobacter crescentus* and 16S ribosomal profiling showed 99.5% homology between the clinical isolate (*Caulobacter sp*. SSI4214) to a common laboratory *C. crescentus* strain CB15 [7]. We performed next-generation Illumina sequencing on the *Caulobacter sp*. SSI4214 strain and created a draft genome assembly to understand the isolate’s relationship to other *Caulobacter* species. Analysis of the 16S rRNA gene obtained from Illumina sequencing confirmed the initial report, with 99.5% similarity to *C. crescentus*. However, phylogenetic reconstruction comparing the 16S sequences of all available whole-genome *Caulobacter* species revealed that *Caulobacter sp*. SSI4214 resides in a separate clade within the *Caulobacter* genus, between *Caulobacter crescentus* and *Caulobacter segnis* (Fig 1A). SSI4214 was also similar to both *C. crescentus* and *C. segnis* with respect to overall GC content and two-way average nucleotide identity (Table 1). Given the convention that isolates of the same bacterial species should have at least 95% average nucleotide identity, our results indicate that SSI4214 represents a distinct species in the *Caulobacter* genus, which we have named *Caulobacter mirare*, as “mirare” is the Latin root for “mirror” and this species mirrors previously-characterized *Caulobacters* (Table 1, S1 Fig) [20].

**Figure 1:**
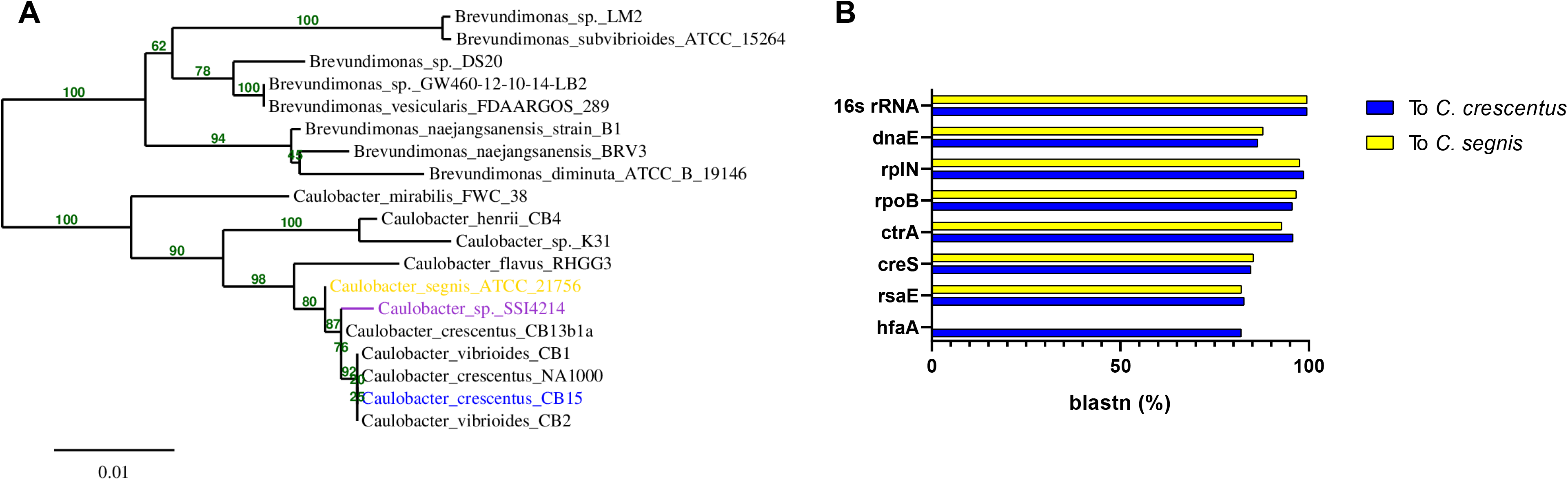
Genomic comparison of *Caulobacter mirare* SSI4214 to related *Caulobacter* species. (A) Phylogenetic tree containing SSI4214 along with closely related species. Numbers indicate bootstrapping confidence values for nodes after 100 replicates. Bar represents average nucleotide substitution/site (B) BLAST values of conserved and *Caulobacter* genus-specific genes.

**Table 1:** Genomic features of *Caulobacter mirare* draft genome assembly compared to *C. crescentus* and *C. segnis*.

|  | *C. crescentus* CB15<br>(complete) | *C. mirare* SSI4214<br>(draft genome) | *C. segnis* TK0059<br>(complete) |
| --- | --- | --- | --- |
| Genome Size (bps) | 4,016,947 | 4,789,750 | 4,655,622 |
| GC Content (%) | 67.21 | 67.51 | 67.67 |
| Predicted-Coding Genes | 3,819 | 4,329 | 4,330 |
| Pathogenicity Islands | 0 | 0 | 0 |
| Average Nucleotide Identity (%) of<br>SSI4214 to | 83.88 |  | 84.75 |

Annotation of the *C. mirare* genome allowed us to compare homology of its genes to those of *C. crescentus* and *C. segnis*, including both broadly-conserved and *Caulobacter*-specific genes [21]. Overall, *C. mirare* is predicted to encode 4,329 protein-encoding genes. This number is similar to that of the *C. segnis* genome (4,330 genes), and larger than *C. crescentus* (3,819) (Table 1) [22, 23]. Among broadly-conserved genes, subunits of DNA polymerase, RNA polymerase, and ribosomes all exhibited at least 86% sequence similarity to both *C. crescentus* and *C. segnis. C. mirare* also possesses clear homologs of many *Caulobacter-specific* genes including the cell-cycle regulator *ctrA*, the curvature determinant *creS*, the S-layer secretion protein *rseE*, and the holdfast attachment protein *hfaA* (Fig 1B). We note that *C. segnis* does not possess a majority of the holdfast synthesis genes, including *hfaA* (Fig 1B) [22]. The observations that *C. mirare* is more similar to *C. segnis* with respect to gene number but more similar to *C. crescentus* with respect to holdfast gene content supports its placement as an independent clade in between the two related species. Importantly, like *C. crescentus* and *C. segnis*, no known annotated virulence factor homologues or pathogen-associated genes are predicted to be present in *C. mirare* [24]. Thus, genome sequencing suggests that the pathogenicity of *C. mirare* is not the result of acquisition of a significant pathogenicity island, and that this clinical isolate broadly resembles environmental *Caulobacter* isolates that were previously considered non-pathogenic.

### Both *C. mirare* and *C. crescentus* are pathogenic towards *Galleria mellonella*

Given the genomic similarity between *C. mirare* and *C. crescentus* we sought to directly compare their pathogenic potential in an *in vivo* host model. *Galleria mellonella*, the greater wax moth, has emerged as a useful system for assessing infection potential due to its relatively short lifespan and ability to inject a defined inoculum of bacteria [25]. Additionally, *Galleria* produces melanin upon infection as part of its immune response, providing a robust visual readout for host health. The process of melanization is irreversible such that even if *Galleria* successfully eliminates the cause of infection, it maintains a dark coloration that corresponds to the degree of its immune response [26, 27].

To quantitatively assay bacterial virulence, we injected *Galleria* with similar numbers of exponentially growing bacteria and monitored melanization after 24 hours, which included fatal events (Fig 2A). As a negative control, we confirmed that mock injections of *Galleria* with water had no effect on melanization. Injection with a lab *E. coli* MG1655 strain also had no effect on *Galleria* melanization, indicating that not all bacteria are pathogenic towards *Galleria* (Fig 2B) [28]. In contrast, injection with *C. mirare* resulted in significant melanization within 24 hours (Fig 2B), suggesting that *Galleria* is a useful model for studying *C. mirare* pathogenesis. Interestingly, injection with two different lab strains of *C. crescentus*, CB15 and NA1000, also resulted in significant *Galleria* melanization within 24 hours (Fig 2B) [29]. The extent of the pathogenesis of the lab *C. crescentus* strains towards *Galleria* was comparable to that of the clinical *C. mirare* strain.

**Figure 2:**
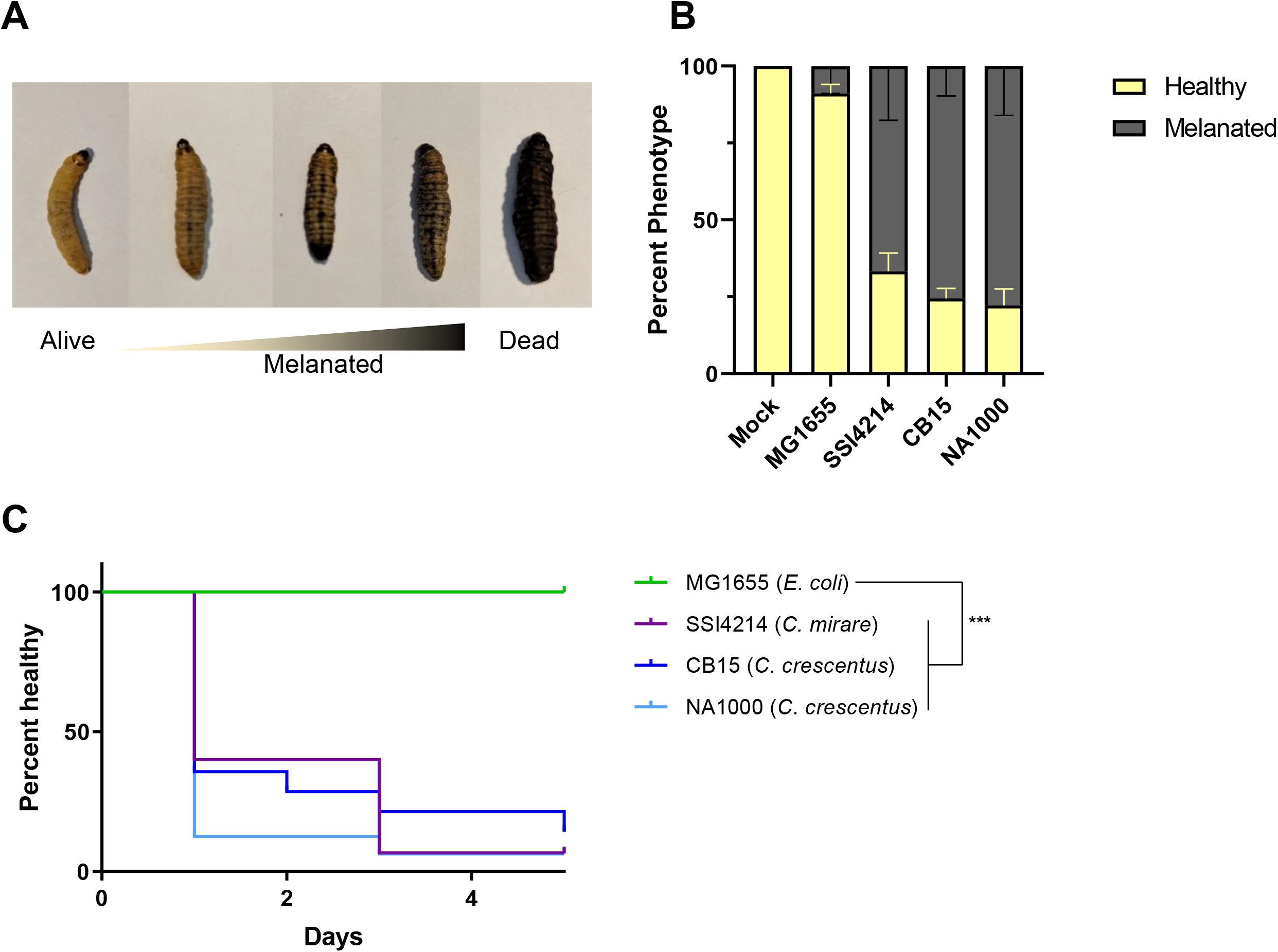
*Galleria mellonella* healthspan decreases upon *Caulobacter* infection. (A) Example images of phenotypes considered for scoring in healthspan assay. (B) Percentage of worms scored as healthy or melanated 24 hours post-inoculation. Error bars represent standard error for three biological replicates. (C) Kaplan-Meier survival analysis for *C. crescentus* strains CB15 and NA1000, *E. coli* strain MG1655 and *C. mirare* strain SSI4214. Survival curve shown is one representative cohort (n=15) of three biological replicates (Mantel-Cox test for statistics, ***P < .001).

To follow the dynamics of pathogenesis we performed a healthspan assay by monitoring melanization as a function of time after injecting *E. coli* (MG1655), *C. mirare* (SSI4214), and *C. crescentus* (CB15 and NA1000). Even five days post injection, no melanization was observed with *E. coli*, validating its use as a non-pathogenic control (Fig 2C). Meanwhile, significant melanization was observed within 1 day of injecting any of the *Caulobacter* strains and increased as a function of time (Fig 2C). *Galleria* injected with the clinical *C. mirare* and lab *C. crescentus* strains displayed similar healthspans (Fig 2C). Together these data suggest that *C. mirare* can function as an opportunistic pathogen, consistent with its clinical isolation and pathology. However, *C. mirare* pathogenesis is not unique, but rather a feature it shares with environmental isolates of *C. crescentus*.

### A cell-associated toxic factor facilitates the pathogenesis of *Caulobacter* in *Galleria*

Since *C. crescentus* infected *Galleria* as well as *C. mirare* but is more experimentally tractable, we focused our efforts on characterizing the mechanism of *Caulobacter* pathogenesis on *C. crescentus*. We first determined whether *Galleria* melanization requires *C. crescentus* growth within the host by heat-killing exponentially-growing bacterial cells prior to injection. Using the same starting number of bacterial cells, heat-killed *C. crescentus* induced similar melanization to living cells (Fig 3A). Thus, the melanization of *Galleria* by *C. crescentus* is not merely a secondary consequence of bacterial growth within the host or outcompeting the host for nutrients.

**Figure 3:**
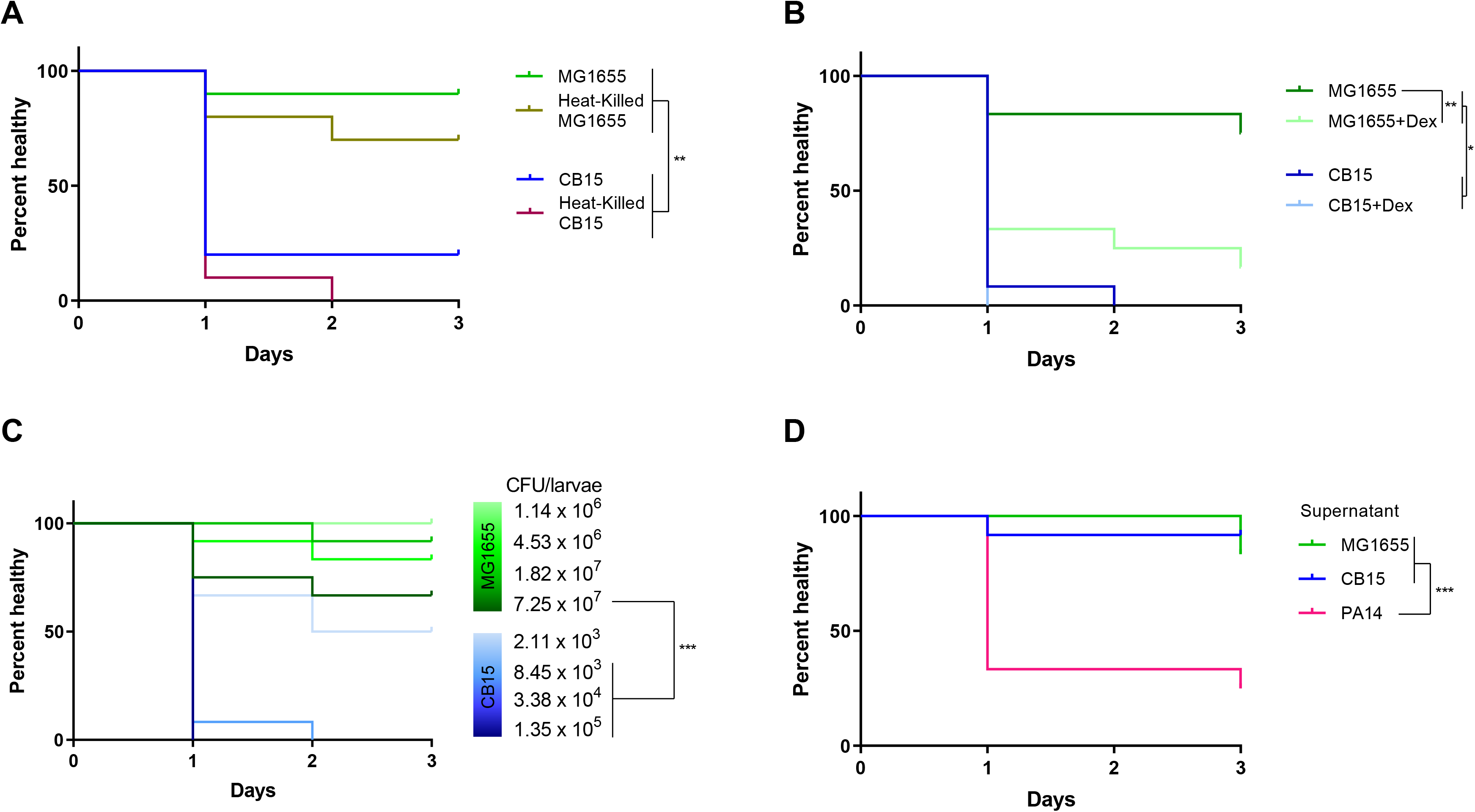
*Caulobacter* pathogenesis is induced by a toxic cell-associated factor. (A) Healthspan of *Galleria* upon injection of 5 μL of OD660 ~ 0.5 (exponentially-growing) live and heat-killed CB15 and MG1655 (B). Healthspan of *Galleria* upon injection with serial dilutions of overnight CB15 or MG1655. (C) Healthspan of *Galleria* upon co-injections of exponentially growing CB15 or MG1655 and 200 ug/larva dexamethasone 21-phosphate (+Dex). (D) Healthspan of *Galleria* upon injection of supernatant derived from overnight cultures of CB15, MG1655, or *Pseudomonas aeruginosa* strain PA14. All survival curves are a representative cohort (n = 10-15) of three biological replicates (Mantel-Cox test for statistics, *P < 0.05, **P < 0.01, ***P < .001).

We next interrogated whether *Galleria* melanization by *C. crescentus* is due to a hyperactivated immune response. Dexamethasone 21-phosphate was previously shown to suppress immune function in *Galleria* by inhibiting macrophage-like haemocyte cells that are responsible for cellular and humoral immunity [30]. Because suppression via dexamethasome would limit activation of the humoral response, decreased melanization upon infection and dexamethasome treatment would indicate that the response is due to immune hyperactivation while increased melanization would indicate that the response is due to bacterial-associated cytotoxicity. We thus co-injected *Galleria* with each of our bacterial strains and dexamethasone. For MG1655 *E. coli* we found that dexamethasone treatment significantly increased melanization (Fig 3B). This finding is consistent with a previous study suggesting that *E. coli* are capable of virulence towards *Galleria* when immunosuppressed by dexamethasone but that the *Galleria* immune system is normally sufficient to prevent *E. coli* pathogenesis [30]. For CB15 *C. crescentus* we found that dexamethasone treatment did not reduce melanization. However, *C. crescentus-induced* melanization is so robust even in the absence of immunosuppression that any such increase is difficult to detect (Fig 3B). These results suggest that active haemocytes prevent *E. coli* from being deleterious in *Galleria* while *C. crescentus* possesses an additional feature that contributes to its ability to infect immunocompetent worms.

Given that the dexamethasone treatment suggested that *C. crescentus* is directly cytotoxic towards *Galleria*, we hypothesized that symptomatic infection is induced via a toxic factor. A hallmark of toxin-associated pathogenesis is quantitative dependence on bacterial load. To assess the bacterial load required to cause an infection phenotype, we injected *Galleria* with four-fold serial dilutions of overnight cultures of CB15 and MG1655 and performed healthspan assays. For both *C. crescentus* and *E. coli*, we observed the expected dose-dependence of infection, with increased melanization as a function of increased numbers of bacteria injected (Fig 3C). This experiment also reinforced the difference in pathogenic potential of the two bacterial species, as the lowest number of *C. crescentus* injected, (~10^3^), caused more melanization than even the highest number of *E. coli* injected (~10^7^).

To determine whether the cytotoxicity of *C. crescentus* is due to a secreted or cell-associated factor, we injected *Galleria* with *C. crescentus-conditioned* media. Specifically, we centrifuged an overnight *C. crescentus* culture capable of inducing *Galleria* melanization at low speeds (5700 g) to remove bacterial cells and cell-associated factors and injected the supernatant that retains secreted factors. CB15 and MG1655 conditioned media did not induce *Galleria* melanization (Fig 3D). As a positive control to confirm that it is possible to induce melanization with secreted toxins we also isolated conditioned media from *Pseudomonas aeruginosa* strain PA14, which is known to secrete exotoxins (Fig 3D) [31]. We confirmed that PA14-conditioned media induced *Galleria* melanization, suggesting that the *C. crescentus* toxic factor is not secreted.

### *Caulobacter* detection in the clinic may be limited due to culturing requirements

*Caulobacters* are ubiquitously present in water systems and our results suggest that they can function as pathogens in some contexts, raising the question of why *Caulobacters* have not been more commonly associated with human infections. One possibility is that *Caulobacter* infections are more common than currently appreciated but that *Caulobacters* are not readily isolated by traditional clinical methods. Because most classical identification methods rely on culturing, we compared the culturing requirements of *C. crescentus* and *C. mirare. C. mirare* was isolated using Danish blood agar plates [7], and we confirmed that the SSI4214 strain indeed grows on sheep’s blood agar (Fig 4A). In contrast, CB15 *C. crescentus* was unable to grow on sheep’s blood agar (Fig 4A). To determine the root cause of this difference, we compared *C. crescentus* and *C. mirare* growth on several complex media. Both species grew robustly on peptone-yeast extract agar, the standard culturing medium for CB15, and nutrient agar. Media with higher salt concentrations, such as Luria broth and terrific broth, did not allow for growth of either *Caulobacter* species. Meanwhile, lower salt-containing media such as tryptic soy agar and super optimal broth, promoted the growth of *C. mirare* but not *C. crescentus* (Fig 4A).

**Figure 4:**
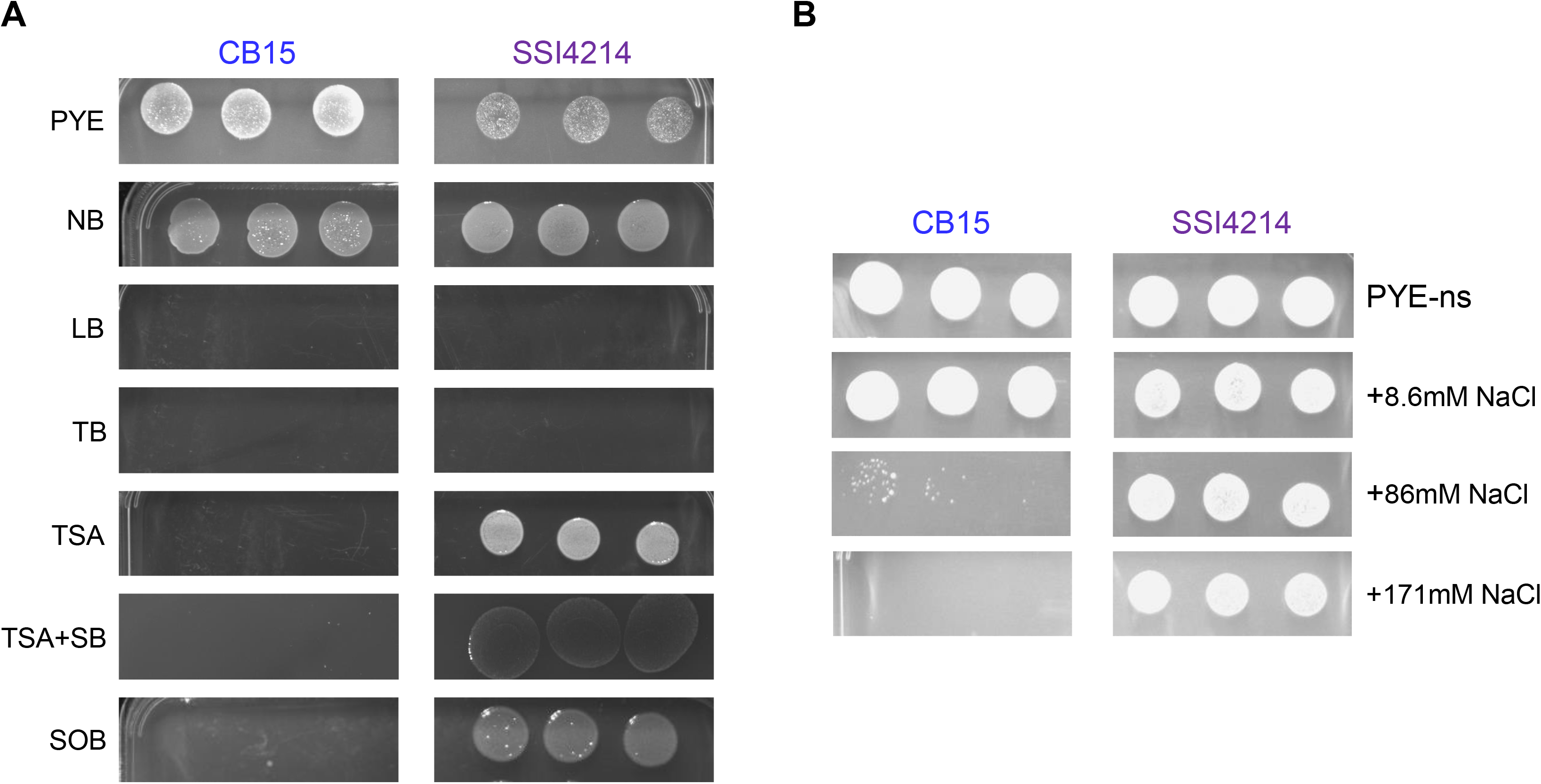
Culturability of *Caulobacter crescentus* (CB15) and *Caulobacter mirare* (SSI4214). (A-B) Three replicates each of 10^−3^-diluted overnight culture of CB15 (left) or SSI4214 (right) on various media. (A) PYE = peptone-yeast extract, NB = nutrient broth, LB = luria broth, TB = terrific broth, TSA = tryptic soy agar, TSA+SB = tryptic soy agar + 5% sheep blood, SOB = super optimal broth. (B) PYE-ns = PYE without added salts, +8.6mM NaCl = PYE-ns with addition of 8.6 mM NaCl, +86mM NaCl = PYE-ns with addition of 86 mM NaCl, +171 mM NaCl = PYE-ns with addition of 171 mM NaCl.

To directly determine if salt content is the relevant growth-determining difference in these media we plated both *Caulobacter* species on PYE in which we replaced the normal MgSO4 salt with varying amounts of NaCl. Both species still grew on modified PYE with no salt added (Fig 4B). We then increased the NaCl content of the modified PYE and found that while both *Caulobacter* species grew well at 8.6 mM NaCl, *C. mirare* continued to grow well at 86 mM and 171 mM NaCl, while *C. crescentus* grew poorly at 86 mM NaCl and failed to grow at all at 171 mM NaCl (Fig 4B). These data suggest that the increased salt tolerance of *C. mirare*, while not necessary for *Galleria* pathogenesis, may explain why this strain could be cultured from an infected patient.

## DISCUSSION

Our work demonstrates that both the clinical *C. mirare* and lab *C. crescentus* species can function as pathogens of *Galleria* with similar degrees of virulence. Not all bacteria can perturb *Galleria* healthspan, as lab strains of *E. coli* proved non-pathogenic in this context (Fig 2). Furthermore, sequencing and analysis of the *C. mirare* genome indicated that this clinical isolate is similar to *C. crescentus* and related *Caulobacters* that were also considered to be non-pathogenic like *C. segnis* (Fig 1). *C. mirare* does not appear to have acquired any clear pathogenicity islands or virulence factors [24]. Coupled with its similar extent of pathogenicity as *C. crescentus*, our findings thus suggest that *C. mirare* is not unique in its ability to cause human disease but that the capacity for virulence may be a general yet previously-unappreciated feature of *Caulobacters*.

In addition to showing that both *C. mirare* and *C. crescentus* can be pathogenic, establishing *Galleria* as a model host for *Caulobacter* provided us with a tractable system for probing the mechanism of pathogenesis. Specifically, we showed that *C. crescentus* pathogenesis persisted in immunocompromised worms, suggesting that *Caulobacter* directly damages the worm as opposed to overactivating the immune system (Fig 3B). Consistent with this finding, all clinical reports of human *Caulobacter* infections occurred in hospital settings with patients who are likely immunocompromised [7–10]. Additional characterization suggested that *C. crescentus* has a cell-associated toxin responsible for its pathogenesis, as *C. crescentus* virulence is dose-dependent, heat-tolerant, and not secreted (Fig 3). One class of cell-associated factors are immunogenic factors such as lipopolysaccharide (LPS) or peptidoglycan (PG) [32, 33]. However, our immunosuppression results suggest that *C. crescentus* pathogenesis is not due to immune hyperactivation and the avirulence of *E. coli* (which has both LPS and PG) indicates that the *C. crescentus* toxic factor is not ubiquitous among Gram-negative bacteria. Alternatively, *C. crescentus* could express a cytotoxic product that actively targets eukaryotic cells. Supporting this model, a survey of bacteria from various aquatic sources demonstrated that *Caulobacter segnis* (the only *Caulobacter* species present in this survey) can directly lyse amoebae [34]. In the future, identification and characterization of the toxin responsible for *Caulobacter* pathogenesis would enable the engineering of *Caulobacter* strains that are less toxic and thus more attractive as vectors for bioengineering or medical applications.

If *Caulobacter* species can generally function as opportunistic pathogens, perhaps even of humans, why is the isolation of *Caulobacters* as human pathogens so rare? Typically, successful pathogens need to survive in the environment of their hosts [35]. *Caulobacter* is often described as an oligotroph since it is found in nutrient-poor environments such as fresh-water lakes and drinking water [3, 36]. However, both our work and previous studies show that *Caulobacter* can also thrive in nutrient-rich culturing conditions (Fig 4). Metabolomic studies of the fluids from common infection sites such as peritoneal fluid, cerebral spinal fluid, and plasma show that these fluids contain metabolites and salt concentrations similar to those in media that support *Caulobacter* growth [37–39]. Thus, it is possible that *Caulobacter* species can grow in human hosts and that the reason they are not often detected is that they are not readily culturable on the media commonly used for clinical microbiology [40, 41]. Consistent with this hypothesis, we showed that *C. mirare* can be cultured on blood agar while *C. crescentus* cannot, likely due to the increased salt tolerance of *C. mirare* (Fig 4). This difference could explain why *C. mirare* could be isolated from a patient and suggests *Caulobacter* infections may be more widespread nosocomial pathogens than previously appreciated. Assessing the true extent of *Caulobacter* as a human pathogen will be aided by implementation of culture-independent pathogen identification method like those based on mass spectrometry or metagenomics [42, 43].

The ability of a classically defined “non-pathogen” like *C. crescentus* to cause disease in the *Galleria* animal model raises the question of what defines a pathogen and are there really non-pathogenic bacteria? Combining our findings with previous work on *C. crescentus* suggests that *Caulobacter* can carry out many of the processes typical of other pathogens, including biofilm formation, antibiotic resistance, killing of non-self bacteria, and toxin production [7, 44, 45]. Unlike the patient-isolated *C. mirare*, the CB15 *C. crescentus* strain studied here is an environmental isolate from a freshwater lake [1]. The ability of this environmental isolate to retain pathogenesis towards an animal host suggests that *Caulobacters* can survive in multiple niches [3, 35]. Both *C. crescentus* and *C. segnis* lack obvious host invasion factors, suggesting that their pathogenesis requires a compromised host and explaining why they are opportunistic pathogens. A recent opinion article suggested that pathogenesis should be viewed as a spectrum and that most bacteria will be pathogenic if they can grow to a sufficient concentration within a host [46]. Our study supports this perspective, suggesting that broadening how we identify and isolate pathogens in clinical settings will allow us to better understand the spectrum of pathogens that actually infect humans. Elucidating the pathogenic potential of more bacteria and the mechanisms by which they cause disease may thus ultimately help combat the challenge of undiagnosed infections.

## MATERIALS AND METHODS

### Bacterial strains and growth conditions

For this study, an overnight culture is defined as a single colony inoculated in 5 ml tubes and grown for 16 hours. Exponential phase cultures were obtained by a 20-fold back dilution of overnight culture in fresh media and grown to an OD_660_ of ~0.5. *Caulobacter crescentus* laboratory strains (CB15 and NA1000) were grown in shaking culture at 30°C in PYE media on platform shakers. *Caulobacter mirare* (SSI4214) was grown in nutrient broth (NB) at 37°C in shaking culture. *E. coli* MG1655 and *Pseudomonas aeruginosa* PA14 were grown at 37°C in LB medium either in shaking culture or roller drum, respectively. Components of organisms’ respective growth media as well as other medias for agar plating have been described previously.

### Genomic analysis and phylogeny construction

Paired-end 150 nt Illumina MiSeq sequencing was performed on all samples at Princeton University’s Genomics Core. Scaffolds were generated from reads using UniCycler default settings on “normal mode,” and assembly metrics were compiled using QUAST [47]. Annotation of the genome was accomplished via DFast with default settings [21]. For phylogenetic construction, an online pipeline (www.phylogeny.fr) was used with default settings. Alignment via MUSCLE was run on “full mode” and phylogeny was determined by bootstrapping with 100 runs. Visualization of the tree was created using TreeDyn [48].

### *Galleria mellonella* healthspan assay

All *Galleria mellonella* larvae were Vita-Bugs© distributed through PetCo© (San Diego, CA) and kept in a 20°C chamber. Larvae were used for healthspan assays within three days of receipt of package. Worms which were not already melanized were assigned randomly to infection or control cohorts. All inoculums were administered using a sterile 1 ml syringe attached to a KD Scientific pump. Same volume injections (5 μL) were delivered at a rate of 250 μl/min to the fourth leg of the worm, which was sterilized with ethanol. Melanization phenotype was determined by observation of a solid black line along the dorsal midline of the larva (Fig 3A). Each figure graph is a representative cohort (n = 10-15 per treatment) from a biological triplicate, except for PA14 which was performed separately (Fig 4D). Mantel-cox statistics for the cohort were calculated using PRISM, and the pooled results are presented in the supplement (S2 Fig).

For co-injection immunosuppression experiments, a dexamethasone 21-phosphate disodium stock solution was made (solubilizing 50 mg/ml in H_2_0) and injected at 200 μg/larva as previously described [29]. Worms not co-injected with dexamethasone-21 were mock injected with sterile H_2_0. For serial dilution experiments, overnight cultures were diluted 4-fold in their respective medium. CFUs were determined by plating overnight cultures on agar plates. For conditioned media experiments, overnight cultures were centrifuged at 5700xg for 3 minutes and the resulting supernatant was injected into the worms. For heat-killing experiments, exponentially growing bacteria were held at 100°C for 10 minutes.

## ACKNOWLEDGEMENTS

We would like to thank Ulrik Justesen (University of Southern Denmark) for sending clinical *Caulobacter sp*. SSI4214 for characterization, and the Princeton University Genomics Core Facility for assistance with genome sequencing. We also would like to thank Gitai lab members Ben Bratton and Robert Scheffler as well was Professor Mohamed Donia for reviewing of the manuscript prior to submission.

