## Supplemental Figures for "Both clinical and environmental *Caulobacter* species act as opportunistic pathogens"

**SUPPLEMENTAL INFORMATION**

Supplemental Figure 1: Average Nucleotide Identity (ANI) plot between *Caulobacter* species

Supplemental Figure 2: Pooled cohort data for healthspan assays

**Supplemental Figure 1: Average Nucleotide Identity (ANI) plot between *Caulobacter* species.** Histogram represents reciprocal best hits (two-way ANI) between fragments of the specified genomes with box-and-whisker plot showing the distribution.

*Caulobacter segnis* ATCC21756 and *Caulobacter mirare* SSI4214

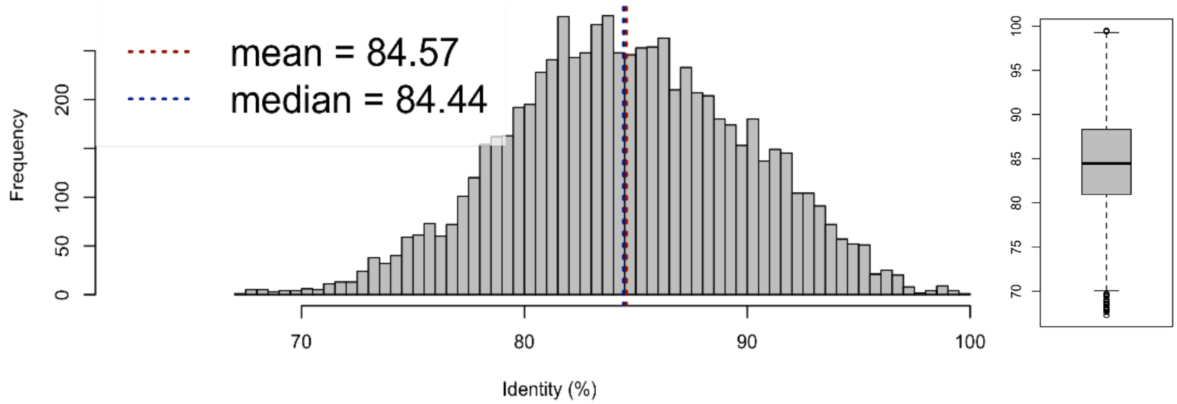


*Caulobacter crescentus* CB15 and *Caulobacter mirare* SSI4214

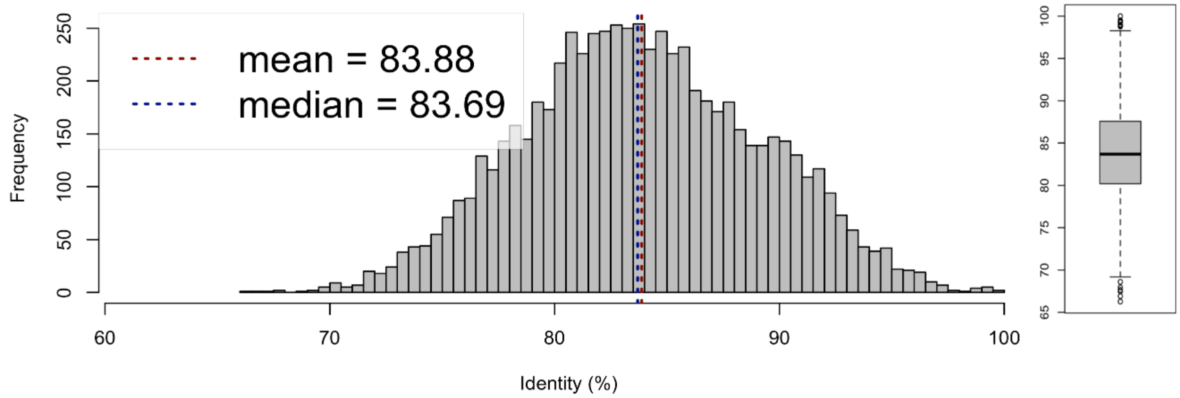


*Caulobacter crescentus* CB15 and *Caulobacter segnis* ATCC21756

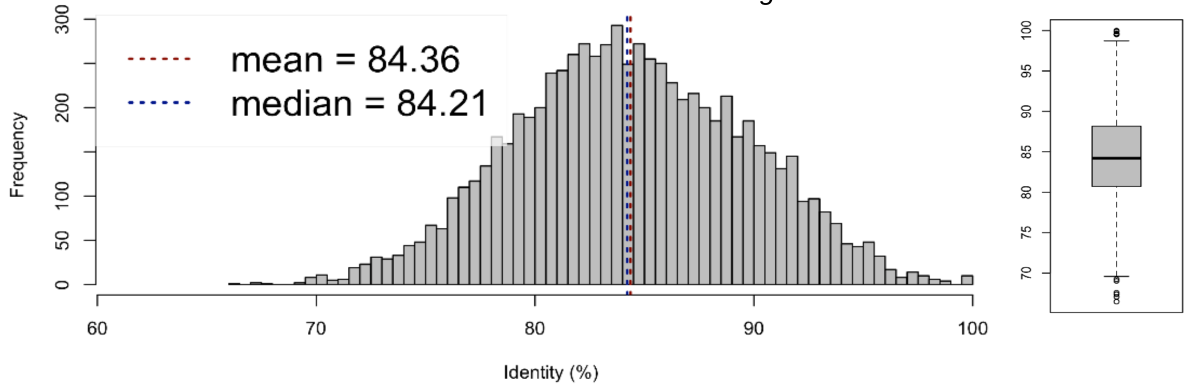

**Supplemental Figure 2: Pooled cohort data for healthspan assay.** Each experiment was performed in biological triplicate. n represents number of animals per cohort and error bars represents standard error.

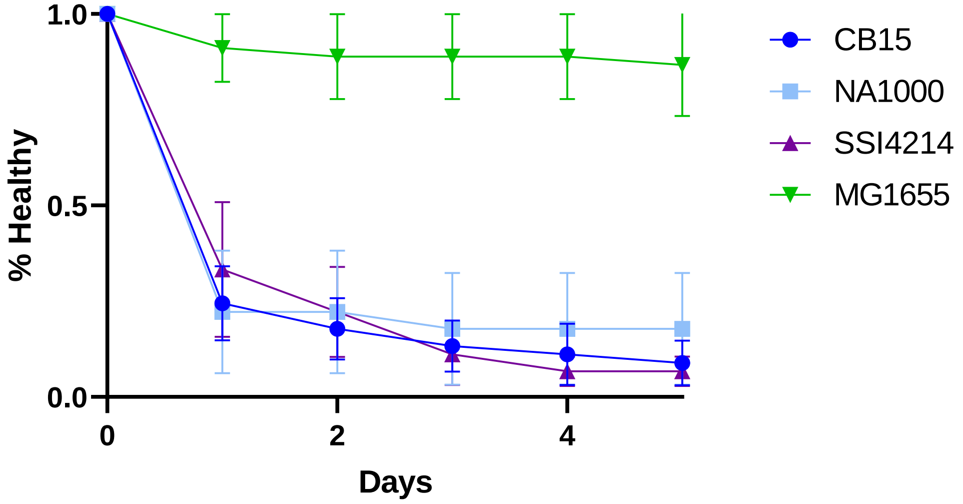


Figure 2C (n = 15)

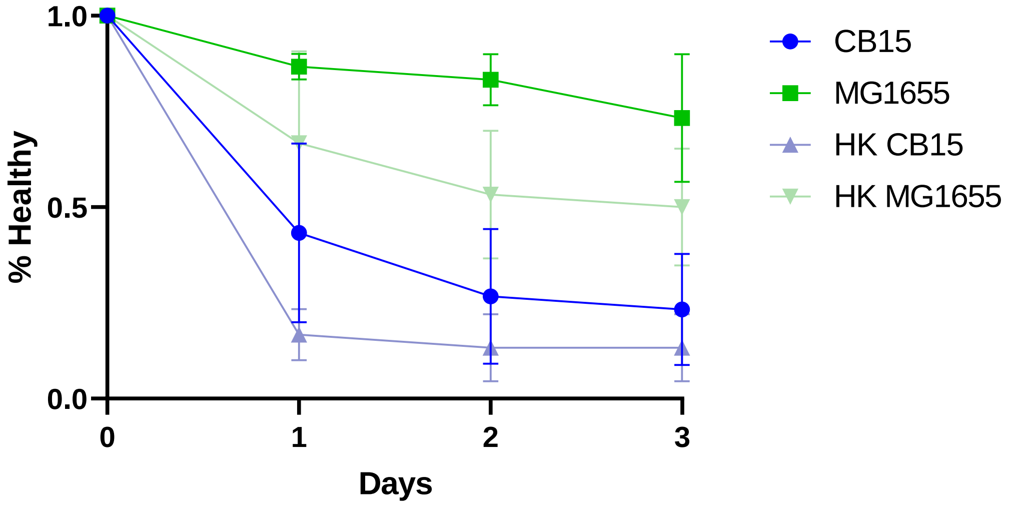


Figure 3A (n = 10)

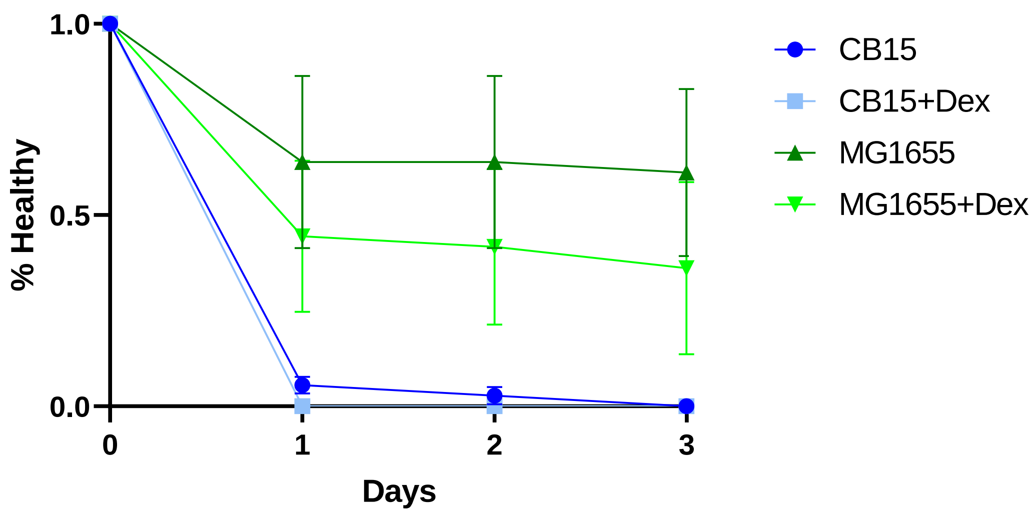


Figure 3B (n = 15)

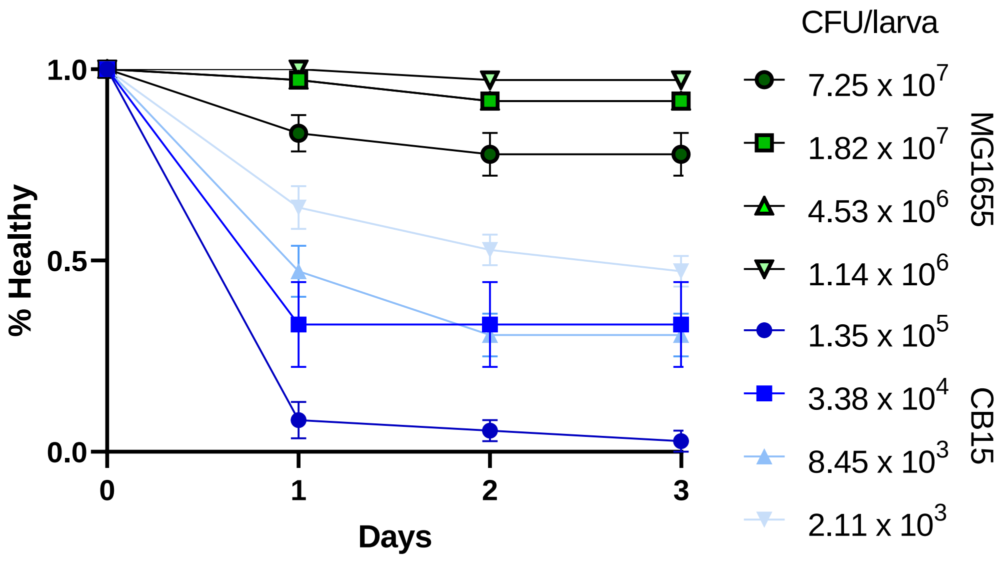


Figure 3C (n = 12)

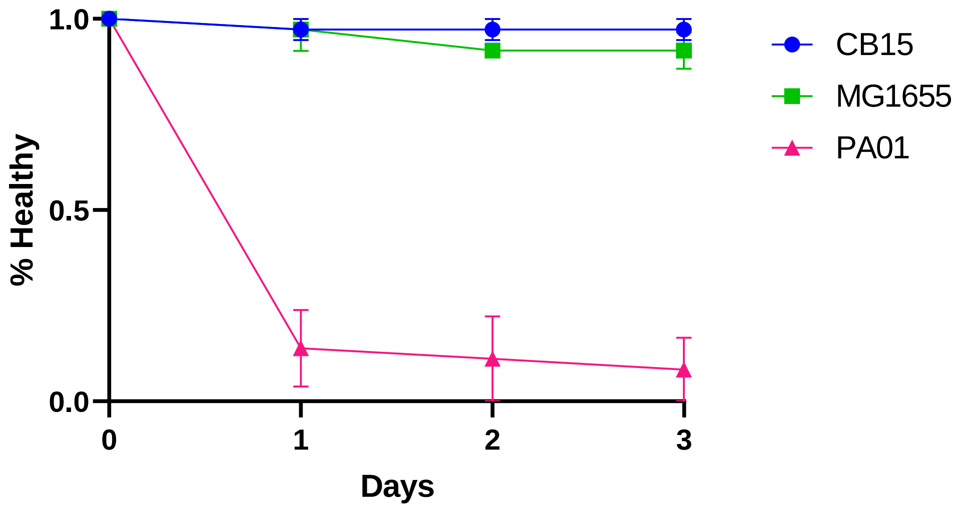


Figure 3D (n = 12)
